# Heterotypic Seeding Generates Mixed Amyloid Polymorphs

**DOI:** 10.1101/2024.03.15.585264

**Authors:** S. Banerjee, D. Baghel, H. O. Edmonds, Ayanjeet Ghosh

## Abstract

Aggregation of the amyloid β (Aβ) peptide into fibrils represents one of the major biochemical pathways underlying the development of Alzheimer’s disease (AD). Extensive studies have been carried out to understand the role of fibrillar seeds on the overall kinetics of amyloid aggregation. However, the precise effect of seeds that are structurally or sequentially different from Aβ on the structure of the resulting amyloid aggregates is yet to be fully understood. In this work, we use nanoscale infrared spectroscopy to probe the spectral facets of individual aggregates formed by aggregating Aβ42 with antiparallel fibrillar seeds of Aβ (16-22) and E22Q Aβ (1-40) Dutch mutant and demonstrate that Aβ can form heterotypic or mixed polymorphs that deviate significantly from its expected parallel cross β structure. We further show that formation of heterotypic aggregates is not limited to coaggregation of Aβ and its isomers, and that the former can form heterotypic fibrils with alpha synuclein and brain protein lysates. These findings highlight the complexity of Aβ aggregation in AD and underscore the need to explore how Aβ interacts with other brain components, which is crucial for developing better therapeutic strategies for AD.

## Introduction

The aggregation of the amyloid β peptide (Aβ) is recognized as a pivotal factor in the pathogenesis of Alzheimer’s disease (AD)^1-4^. Aβ40 and Aβ42, comprising 40 and 42 residues respectively, constitute the primary components of amyloid plaques found in the brains of AD patients^2,4,5^. The aggregation pathway of these peptides involves the spontaneous assembly of misfolded peptides into oligomers, which act as nuclei, and facilitate the formation of protofibrils and ultimately mature fibrils with a characteristic cross-β secondary structure. This aggregation, as studied in-vitro, is a stochastic process influenced by various factors such as concentration of monomers and/or oligomers, temperature, pH, nature of counterions and many more^6-9^. Introducing aggregated species of the same peptide into the aggregation mixture as seeds can also modulate the kinetics of aggregation^10-13^. Several studies have demonstrated that the presence of seeds can shorten the lag phase of the aggregation process, with the seed surface serving as a template for generating new aggregates through secondary nucleation. Such templated growth is expected to retain the structural form of the seed, and as a result is commonly used to investigate the structure of fibrils derived from brain protein extracts of AD patients^13-15^. While the effect of homotypic interactions leading to seeded growth of fibrils is well understood, one key aspect of amyloid aggregation that has perhaps been somewhat under investigated is the effect of heterotypic interactions. Essentially, it is not well understood how the presence of a fibrillar seed that is sequentially and/or structurally different, i.e., heterotypic to Aβ can modulate the aggregation process. Cukalevski et al. have investigated the coaggregation of Aβ40 and Aβ42 and identified evidence of interactions between the peptides at early stages of aggregation, however these intermediates ultimately do not lead to formation of mixed or heterotypic fibrils at later stages^16^. These findings have also been validated by Baldassarre and coworkers, who have demonstrated the formation of mixed oligomers through coaggregation of the peptides^17^. Yoo and coworkers have demonstrated that the fibrillation of the Arctic mutant of Aβ40 can be seeded by Aβ42, but not of Aβ40^18^. Pansieri et al. have demonstrated templating of S100A9, a proinflammatory protein involved in AD, on the surface of Aβ42 amyloid fibrils^19^; however, the opposite effect, i.e., how a different protein can affect the aggregation pathway of Aβ, is yet to be fully understood. Furthermore, while Aβ40 and Aβ42 are known to be the major isoforms of Aβ in the context of AD, amyloid plaques from AD brain specimens have been shown to contain other N- and C-terminal truncated isoforms of Aβ as well^20,21^. In general, the interaction between different isoforms of Aβ and how that affects the corresponding aggregation pathways has been subject of very limited number of studies. Braun and coworkers have shown that the C-terminal truncated Aβ37 and Aβ38 slow down the kinetics of fibrillation of Aβ42, but this is not mediated through formation of heterotypic aggregates^22^. Shorter isoforms of Aβ, particularly those encompassing the amyloidogenic domain between residues 16-21, can adopt a different structural arrangement in the fibrillar phase, namely antiparallel β sheets, as has been conclusively demonstrated^23,24^. However, how such an antiparallel seed can modulate the aggregation of Aβ40 and Aβ42, both of which are known to form fibrils with parallel β structure, is not known. The aggregation of Aβ can involve transient intermediate fibrillar aggregates that have an antiparallel structural motif; presence of such intermediates has been unequivocally demonstrated for the Iowa mutant (D23N) and is speculated to exist for wild type (WT) Aβ as well^12,25^. Fibrils derived from Aβ aggregates in Cerebral Amyloid Angiopathy (CAA), a neuropathy sharing significant overlap with AD, have been shown to contain both parallel and antiparallel motifs^26^. Recent cryo-EM structures of AD brain derived fibrils have also come to similar conclusions^15^. Taken together, these findings point to possible scenarios wherein interactions between Aβ aggregates of different structural arrangements are likely; however, the effect of such interactions remains unclear. All of the above allude to some key questions regarding the mechanism of Aβ aggregation, namely whether it can be seeded with heterotypic fibrillar seeds, and if so, whether the structural motif of the seed propagates in downstream aggregates. Of course, this is not limited in scope to aggregation of Aβ in isolation. Amyloid plaques have been shown to contain proteins such as tau and alpha synuclein in addition to Aβ^20,21,27^; yet how the coaggregation/cross-seeding with these proteins can alter the aggregation pathway of Aβ remains to be fully understood. The interest in understanding heterotypic amyloid interactions have grown significantly in recent years, leading to key insights into how they may affect the aggregation process^19,22,28-32^. The interactions of Aβ with tau have been investigated, and it is known that tau inhibits maturation of Aβ oligomers to the fibrillar stage^33^. However, the generalizability of this finding to interacts of Aβ with other plaque associated proteins is debatable. Furthermore, the precise nature of the interaction of the two proteins that leads to modulation of Aβ aggregation, specifically if it is mediated through formation of mixed/heterotypic aggregates, is not understood. It is unknown if Aβ can form mixed aggregates in presence of other proteins/peptides, either structurally or sequentially heterotypic, or both. While there is growing need to better understand the effect of protein-protein interactions in modulating amyloid aggregation pathways, investigating heterogeneous mixtures of different proteins/peptides presents a major experimental challenge. Conventional biophysical techniques used for amyloid structural elucidation, such as Nuclear Magnetic Resonance (NMR) spectroscopy, can identify interactions between different aggregates and the presence of mixed/heterotypic fibrils^7, 13,32,34^; however, this becomes significantly more challenging when multiple structurally different species are present, since these approaches cannot probe the spectral facets of individual aggregates. Cryo-Electron Microscopy has provided an alternative to deducing fibril structures with atomic level detail^15,35-38^; however, this also remains limited to largely homogeneous structural ensembles. As a result, these approaches have been mostly focused on characterizing mature fibrils, which correspond to a narrowed and homogenized ensemble, but not always towards interrogating the transient structures that lead to fibrils. Hence, the majority of the studies on heterotypic interactions between amyloid proteins have focused on alterations in morphology and aggregation kinetics, but not necessarily on the structure of the aggregates. Recent technological advances in vibrational spectroscopic imaging have opened up possibilities towards probing the spectral parameters of individual aggregates by integrating infrared (IR) spectroscopy with Atomic Force Microscopy (AFM). AFM-IR utilizes the photothermal response of the specimen and its modulation of the AFM cantilever oscillations upon resonant IR excitation, resulting in the measurement proportional to the IR absorbance with nanoscale spatial resolution^39-41^. AFM-IR is thus ideally suited for characterization of heterogeneous amyloid aggregates in dynamic equilibrium as it reports on the spectral and hence structural characteristics of specific members of the structural ensemble, which is beyond the capabilities of spatially averaged bulk spectroscopies. In this report, we leverage the unique capabilities of AFM-IR to demonstrate for the very first time that when incubated with structurally heterotypic antiparallel fibrillar seeds of Aβ (16-22) and Dutch mutant of Aβ-40 (E22Q), Aβ42 forms structurally unique mixed fibrillar aggregates. Seeding with Aβ(16-22) results in two distinct fibrillar polymorphs, one of which is of mixed composition, while the other primarily constitutes of only Aβ42. Interestingly, this ‘pure’ polymorph is structurally different compared to fibrils of Aβ42 formed via unseeded aggregation. Seeding with the Dutch mutant leads to formation of a single polymorph that is comprised of both seed and Aβ42. Furthermore, Aβ42 aggregated in presence of alpha synuclein and brain protein lysates without any fibrillar seeds also forms structurally heterotypic fibrils, which suggests the plausible interaction of Aβ42 with other brain proteins and underscores the generality of our findings. These results underscore the necessity to investigate the structural evolution of amyloid peptides in presence of other partner amyloids for a granular understanding of disease mechanisms.

## Results and discussion

To understand how the structure of the Aβ aggregates is modulated in presence of different seeds, we have used ^13^C-labeled Aβ42 as our reference system. The isotopic substitution of Aβ42 is necessary to spectrally isolate its signal from other proteins/peptides present in the aggregation mixture. To establish the structural benchmarks of pure ^13^C Aβ42, we first performed a control experiment where Aβ42 was allowed to aggregate in absence of any seed. The details of the aggregation procedure are provided in the Supporting Information. AFM images of Aβ42 aggregates after 6 hours of incubation shows the presence of fibrillar aggregates (Figure 1A). Representative IR spectra in the amide I region, obtained from different spatial locations on the fibrillar aggregates, are shown in Figure 1B. The spectra exhibit an amide I peak at ∼1590-1600 cm^-1^ with a shoulder at 1626 cm^-1^ (Figure 1C). Since Aβ42 peptide backbone in our experiment is uniformly ^13^C labeled, the amide frequency undergoes a downshift of ∼30 cm^-1^ compared to unlabeled Aβ42^42,43^. Hence a peak at ∼1590-1600 cm^-1^ denotes that β-sheets are the primary structural component in these aggregates, while the shoulder at 1626 cm^-1^ can be attributed to disordered and/or β-turn structural motifs. The spectra are, however, not identical and exhibit significant variations. After 24 hours of incubation, the spectral heterogeneity is still observed from fibrillar aggregates, and the spectra show similar features to that of 6 h (Figure 1D-F). The spatial variations in the spectra indicate a heterogeneous structural ensemble, which has been previously observed for wild-type Aβ42 and are thus consistent with prior structural studies of Aβ42. These measurements serve as the benchmark for the assessment of the modulation of Aβ42 aggregation by heterotypic seeds.

**Figure 1.**
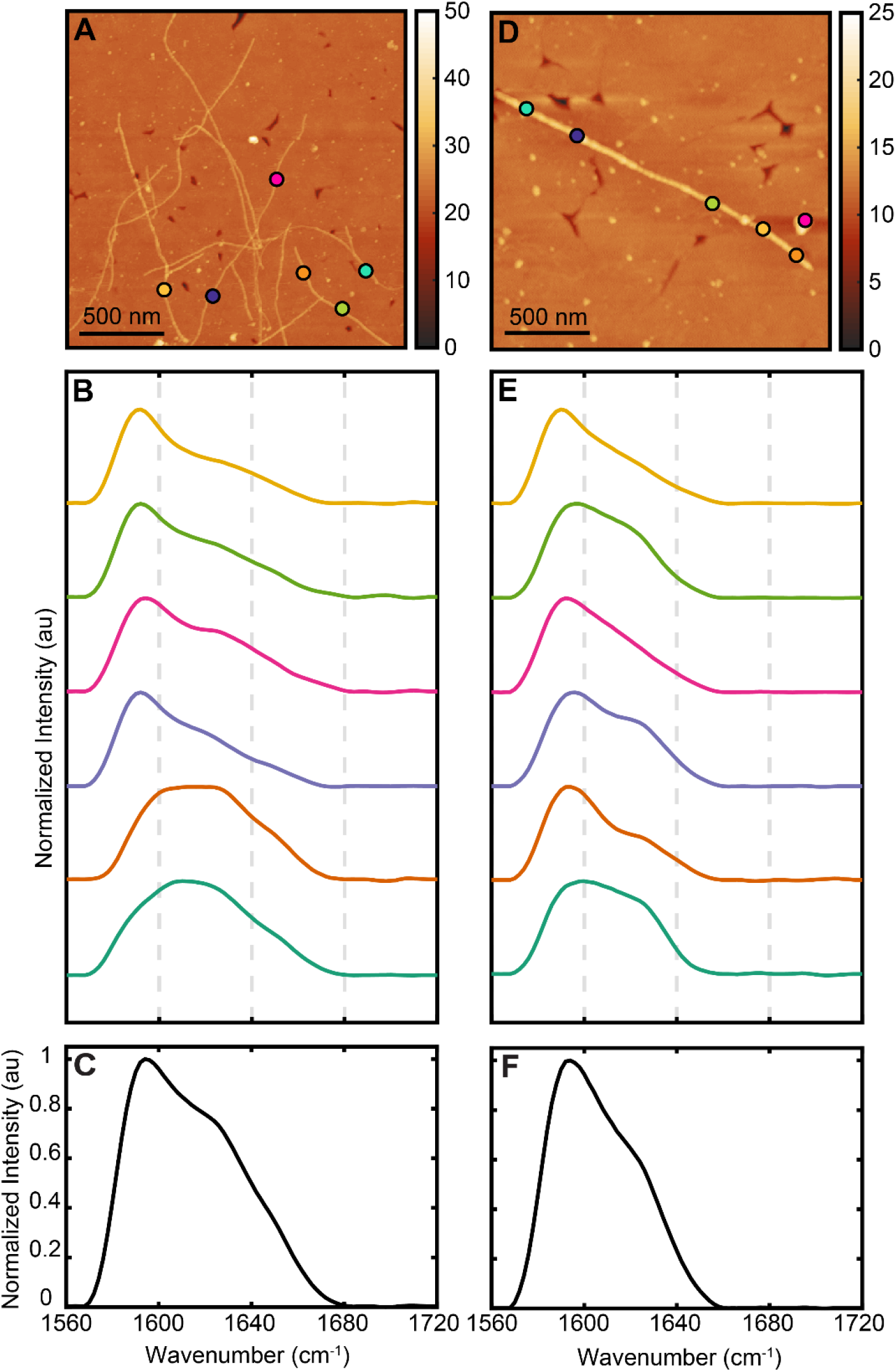
(A,D) AFM image of ^13^C-Aβ-42 fibrils after 6 h and 24 h of incubation at 37 °C, without agitation. (B,E) Representative IR spectrum of amide 1 region recorded from ^13^C-Aβ-42 fibrils and (C,F) Average IR spectrum from the corresponding representative spectra.

The first peptide we chose as seed was Aβ(16-22), which is a small fragment of full length Aβ and corresponds to one of its most amyloidogenic sequences. Aβ(16-22) is well-known to form ordered fibrils with antiparallel arrangement of strands. AFM morphological maps of the Aβ(16-22) fibrils used as seed and representative IR spectra are shown in Figure S1. The spectra exhibit an amide I band with two distinct peaks at 1626 cm^-1^ and 1690 cm^-1^. For unlabeled peptides, an amide I peak at ∼1630 cm^-1^ demonstrates the presence of β-structure and the strong band at 1690 cm^-1^ confirms the anti-parallel nature of β-sheet arrangement. The Aβ(16-22) fibrils represent a sequentially homotypic but structurally heterotypic seed for Aβ aggregation: while the sequence of the seed matches exactly with a segment of the full-length peptide, the secondary structure of the seed is different from those adopted by full-length Aβ. After 6 h of seeded aggregation, two distinct polymorphs can be observed (Figure 2 A,D). The first corresponds to tape like flat fibrils having larger width (102.2 ± 10.7 nm) (Figure 2A), similar to those observed for the seed, while the second exhibits a more rounded morphology with lower width values (20.7 ± 0.7 nm) (Figure 2D), akin to the pure Aβ fibrils. The fibrillar morphology would suggest that the ensemble essentially separates into seed and Aβ aggregates; however, IR spectra from the fibrils portrays a different picture. IR spectra recorded at several points on the seed-like polymorphs, shown in Figure 2B-C, exhibits two peaks at 1628 cm^-1^ and 1692 cm^-1^ similar to Aβ(16-22), with the exception of an additional shoulder that is present at 1600 cm^-1^. The shoulder is conspicuously absent in the seed spectra (Figure S1), but is evident in spectra of ^13^C Aβ42 (Figure 1B-C), thus indicating the mixed or heterotypic nature of these fibrils, which are primarily composed of the seed Aβ(16-22) but also concurrently contain ^13^C Aβ. The other polymorph, that morphologically resembles fibrils of pure Aβ, exhibits an amide I peak at 1628 cm^-1^, and lacks the characteristic peak at 1690 cm^-1^ of the antiparallel seed, indicating these fibrils are mainly composed of Aβ42 (Figure 2E-F). However, the spectra are not identical to those acquired from pure Aβ42 fibrils, where the amide I peak was at 1600 cm^-1^, indicating a different secondary structure of fibrils resulting from seeded growth. The same polymorphs are observed after 24 h of aggregation as well (Figure 2 G, J), and the spectra of each fibrillar morphology are very similar to those observed after 6h (Figure 2 H-L). Additional spectra and AFM images for both timepoints are shown in Figures S2 and S3 respectively. Taken together, these results evidence formation of structurally distinct polymorphs due to seeded aggregation of Aβ42. The invariance of the polymorphic ensemble also suggests that these aggregates resulting from coaggregation are stable and likely not transient intermediate structures that might eventually evolve into a different structure. Interestingly, no fibrils were found that were spectrally identical to exclusively Aβ(16-22), indicating that both Aβ42 and seed interact with each other and incorporate themselves into a single fibril, when they are allowed to aggregate in a mixture.

**Figure 2.**
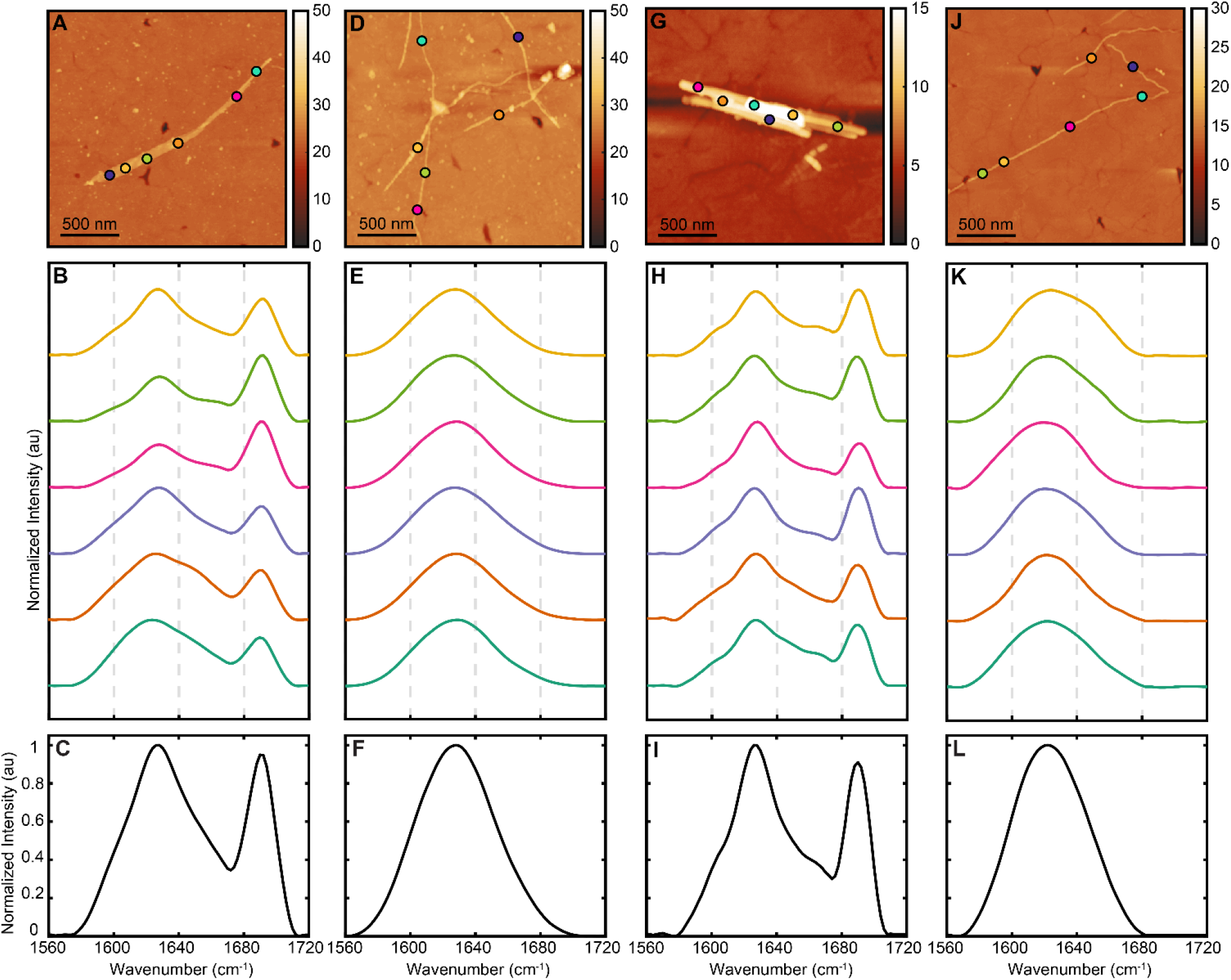
AFM-IR characterization of ^13^C-Aβ42, cross-seeded with Aβ(16-22) fibrils in 10 mM phosphate buffer. (A,D) AFM topographic images of fibrils after 6 h and (G,J) after 24 h of incubation. (B,E,H,K) Representative IR spectrum of cross-aggregates recorded from the corresponding AFM images, where (B,H) demonstrates first spectral subtype coming from flat fibrils, while (E,K) shows the second spectral subtype of round fibrils. (C,F,I,L) represents average IR spectra from the corresponding representative spectra.

To verify if the modulation of Aβ42 aggregation by structurally heterotypic seeds is not limited to specifically Aβ(16-22), we performed the cross-seeding experiments using a different fibrillar seed, namely the Aβ Dutch mutant. The Dutch variant of Aβ differs from the wild type through a point mutation (E22Q)^44^. However, from a structural perspective, the Dutch mutant deviates significantly from wild type Aβ, and has been shown to form stable fibrils exhibiting concurrent parallel and antiparallel character, which offers two structural motifs for seeding^45,46^. The AFM topography and representative IR spectra of the Dutch mutant seed, shown in Figure S1 are consistent with previous reports. Essentially, the spectra lack a pronounced peak at ∼1630 cm^-1^ indicating that the structure of the Dutch mutant seeds is not primarily composed of parallel β structure. The results from seeded growth of ^13^C Aβ after 6h are shown in Figure 3 A-D. Unlike Aβ(16-22), the Dutch mutant leads to a single polymorph; however, we still observe two spectral subtypes (Figure 3B); one with a peak at ∼ 1598 cm^-1^ with a distinct shoulder at 1636 cm^-1^, while the other type exhibits a peak at ∼ 1632 cm^-1^ without a distinct shoulder at ∼1598 cm^-1^. The spectral variations arise from the same fibrillar aggregates, which points to a fundamental difference in how Aβ structure is modulated by the Dutch mutant seed. Unlike Aβ(16-22), which lead to two distinct polymorphs, we get a single polymorph which is structurally heterogeneous, as evidenced by the spectral variations. After 24 h of aggregation, we observe essentially the same ensemble: the AFM images still exhibit only one fibrillar morphology (Figure 3E), and the spectra, shown in Figures 3F-H, still correspond to the two types observed earlier. Additional spectra are shown in Figure S4. While at first glance this may appear to be similar to the aggregation of pure Aβ, the spectral subtypes do not appear identical to the unmodulated control Aβ fibrils, which indicates some effect of the seed.

**Figure 3.**
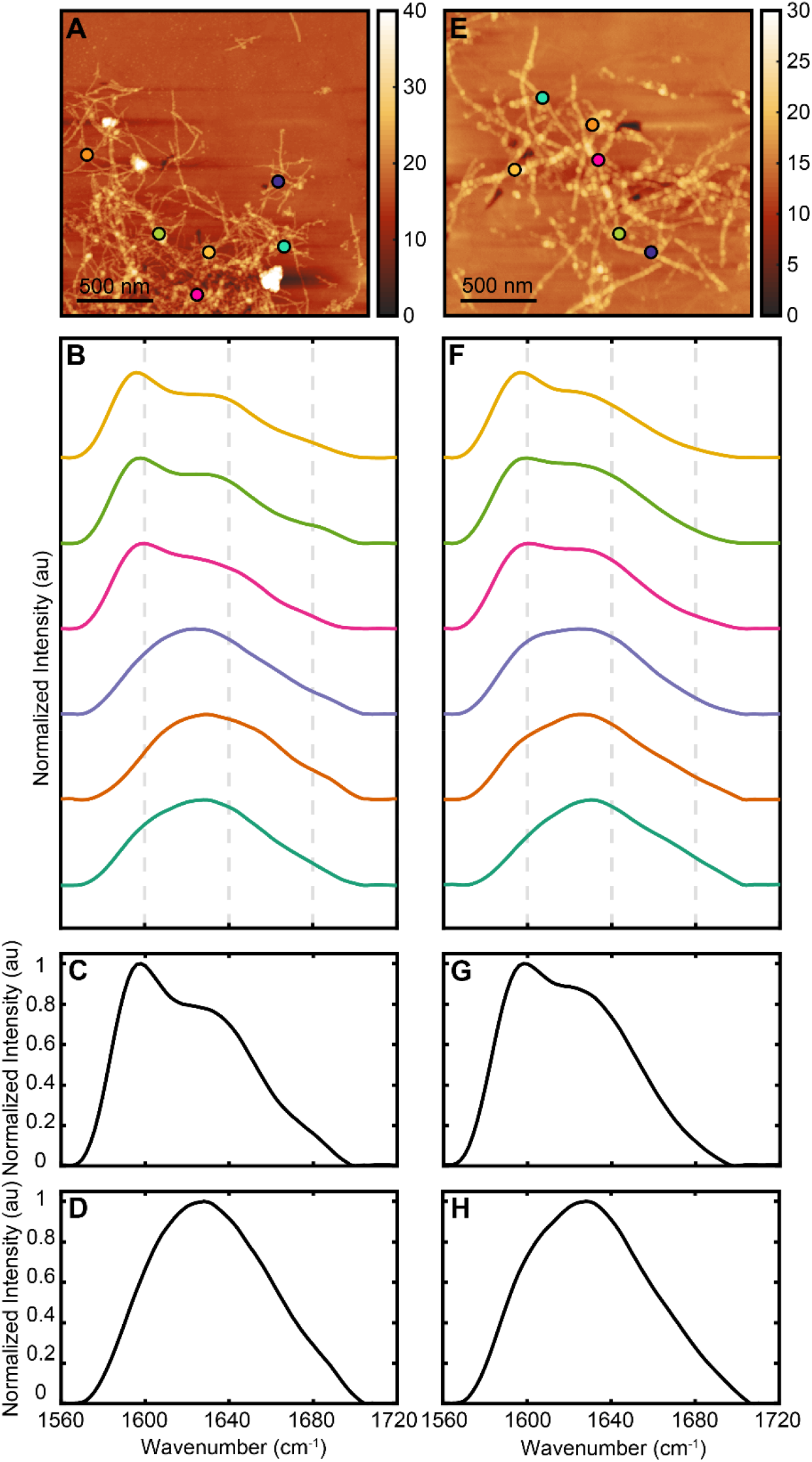
AFM-IR characterization of ^13^C-Aβ42, cross-seeded with fibrils of Dutch Aβ-40 in 10 mM phosphate buffer. (A,E) AFM topographic images of fibrils after 6 h and 24 h of incubation, respectively. (B,F) Representative IR spectrum of cross-aggregates recorded from the corresponding AFM images demonstrating the two spectral subtypes. (C,D,G,H) represents average spectra from the corresponding representative spectra.

However, a detailed understanding of the secondary structure of the above Aβ fibrillar aggregates generated by seeded growth and their putative is not possible without further analysis, this is particularly necessitated by the distinct spectral subtypes observed for the control and seeded fibrils. The characteristic absorption peak for β sheets in unlabeled peptides, parallel or antiparallel, appears at ∼1620-1630 cm^-1^, whereas an additional peak at ∼1690 cm^-1^ is present only in antiparallel structures. This is in excellent agreement with the Aβ (16-22) seed spectra. Non β structural components, such as random coils, turns etc. absorb at ∼1650-1670 cm^-1^. In ^13^C labeled peptides all of these bands redshift by ∼30 cm^-1^. In presence of both labeled and unlabeled peptide, as observed here, the spectral bands from different elements overlap, making unequivocal assessment of structure difficult. Therefore, to gain more quantitative insights into the structural distribution of the fibrils, and elucidate their heterotypic/mixed nature, we deconvoluted the IR spectra using the MCR-ALS algorithm. MCR-ALS is a spectral deconvolution approach akin to spectral global fitting and Singular Value Decomposition (SVD), wherein each spectrum is approximated as a weighted linear combination of a set of basis spectra^47-49^. The basis spectra correspond to specific secondary structural elements in this context, and spectral variations can be interpreted in terms of changes in their relative weights. It should be noted that spectral fitting is the gold standard of deconvolution approaches but requires additional knowledge regarding the number and nature of constituting bands. Details of the MCR-ALS approach are provided in the Supporting Information. For clarity, we deconvoluted the spectra as a superposition of 4 components, corresponding to the 4 main peaks identified in the spectra, as shown in Figure 4A. The component at 1592 cm^-1^ can be attributed to ^13^C Aβ42 with β sheet structure, while the one with peaks at 1690 cm^-1^ and 1626 cm^-1^ corresponds to the unlabeled Aβ(16-22) seed. We also observe a component centered at 1624 cm^-1^ without any high frequency component, and therefore also likely corresponds to primarily ^13^C Aβ42 with disordered/β turn structure. The fourth component is peaked at 1660 cm^-1^. This can be nominally attributed to be representative of non-β sheet structures in the seed. Since different spectral types were observed for some of the polymorphs, we used the weights obtained through MCR to categorize the spectra using k-means clustering. Since k-means is an unsupervised clustering algorithm, the spectral subgrouping is devoid of any biases. This allows for better comparison of the secondary structural distributions between the different polymorphs observed. The mean weights for each spectral class are shown in Figure 4B, which clearly demonstrate the structural differences between pure Aβ and the seeded polymorphs. For clarity, the composition of the two different seed fibrils, is also shown in Figure 4B. We see that the two subtypes of the control Aβ fibrils differ primarily with respect to the relative amount of β sheet and random coil components. The mean weights of spectral components obtained from two seeded fibrillar polymorphs are markedly different in terms of composition from both the control and seed fibrils. One is more similar to the seed, but noticeably contains contributions from the labeled β sheet component, which can only arise from coaggregating ^13^C Aβ. This confirms the heterotypic nature of this category of fibrils. The other polymorph is harder to uniquely attribute as a heterotypic fibril: we do not observe significant contribution from the seed spectra, specifically of the antiparallel component. However, the secondary structural distribution of this group is nonetheless significantly different from pure Aβ fibrils of either type, indicating that these fibrils are structurally distinct from the control aggregates and represent a new polymorph resulting from seeded growth. Interestingly, the key spectral difference seeded fibrils is the relative increase in the 1660 cm^-1^ component. In unlabeled peptides, antiparallel β structures exhibit a characteristic band at 1690 cm^-1^. We observe this band in the seed spectra. Upon isotope labeling, this band is expected to redshift by ∼30 cm^-1^, and thus would appear at ∼1660 cm^-1^. An increase in this component in the seeded fibrils thus suggests enhanced antiparallel character. Hence, the picture that emerges from this analysis is that cross seeding of Aβ with antiparallel fibrils modulates the aggregates and leads to formation of polymorphs that adopt the antiparallel character of the seed. This is a unique result and to the best of our knowledge has never been directly demonstrated before. Essentially, this implies that heterotypic seeding of Aβ is possible, and hence offers a new perspective into aberrant aggregate structures that have been identified in course of Aβ aggregation. For the fibrils seeded by the Dutch mutant, only one fibrillar polymorph is observed which exhibits spectral variations akin to pure Aβ. While this would point towards an unperturbed aggregation mechanism with minimal effect of the seed, the structural composition of the two spectral subtypes differ significantly from pure Aβ. The Dutch mutant seed is composed largely from the components corresponding to random coils and antiparallel β sheets. However, the fibrils seeded by the Dutch mutant exhibit all four spectral components, which indicates integration of ^13^C Aβ and thus unequivocally highlights their heterotypic composition. Both seeded spectral types also exhibit enhanced contribution from the 1660 cm^-1^ component compared to the control, which further validates the antiparallel arrangement in these fibrils. We discuss the implication of these results below.

**Figure 4.**
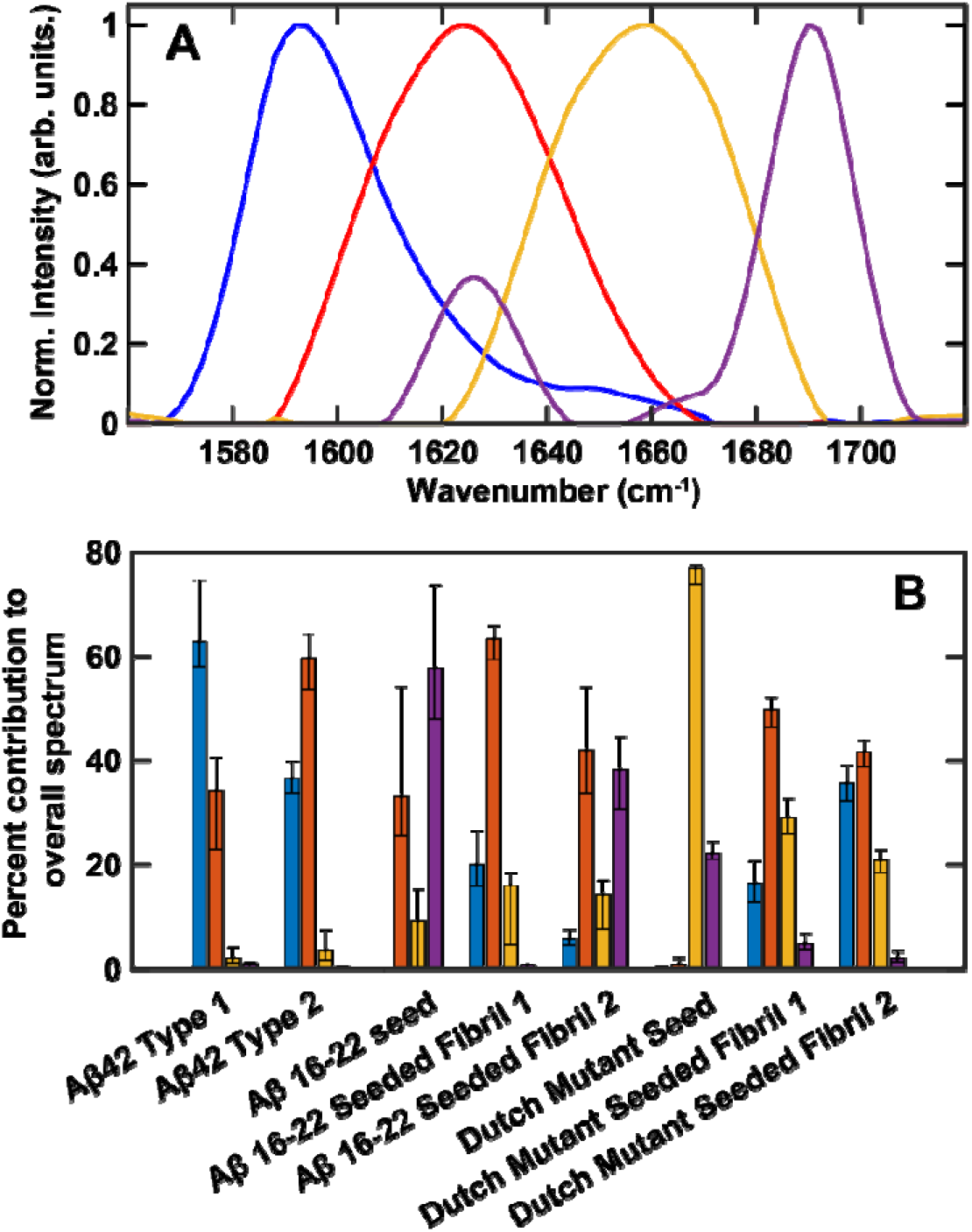
Spectral deconvolution of seeded and unseeded Aβ42 fibrils using MCR-ALS. (A) Components constituting the fibrillar spectra, as determined from MCR (B) Mean percentage weights of each spectral component for different fibrillar aggregates, which is reflective of the secondary structural distribution. The bars are color-matched to the spectral bands in A. The error bars represent the interquartile range of the calculated weights from MCR.

The above observations, taken together, reveal unique mechanistic insights into amyloid aggregation. A core tenet of amyloid aggregation mechanisms is morphological preservation, i.e., a specific parent polymorph will seed morphologically identical filial generations^13, 25^. However, we observe that this does not necessarily hold true for heterotypic aggregation. A single distinct Aβ (16-22) polymorph leads to formation of morphologically divergent daughter fibrils that either resemble the seed or those formed by aggregation of pure Aβ. Seeding with the Dutch mutant, however, leads to only one polymorph and preserves the morphology of the seed in daughter fibrils. Another key aspect of amyloid aggregation is structural retention of parent fibrils in later generations in addition to morphology. This is not just limited to seeding of parallel cross β structures; Tycko and coworkers have demonstrated that antiparallel fibrils of the Iowa mutant of Aβ can seed offspring fibrils of the structural arrangement^12, 25^. This has also been validated in more recent studies involving the Dutch mutant as well^45, 46^. Our observations are consistent with this, as we observe antiparallel character in the mixed fibrils. The key factor implicit in all of the above is the sequence homology of the seed and Aβ. Essentially, when seeded by different polymorphs of the same protein, the daughter fibrils can retain the structural identity of the parent, even if that differs from its natural aggregation state. However, there are increasing number of reports that evidence the existence of heterotypic interactions in amyloid aggregation and their potential impact on aggregation kinetics and toxicity. For example, structures of mixed fibrils of synuclein and the TAR DNA-binding protein (TDP-43)^32^, and a heterotypic amyloid signaling complex^29^ have recently been identified by NMR. It has been shown that interactions with homologous non-Aβ peptides and different isoforms can alter aggregation kinetics and fibril morphology of Aβ^22, 28^. This also applies to other amyloidogenic proteins. Different proteins with sequence homology to the amyloidogenic region of tau have been demonstrated to alter its aggregation pathway^30^. This suggests seeds from different Aβ isoforms may structurally alter WT Aβ aggregates through heterotypic interactions, and not just their rate of formation. However, this has never been specifically evaluated. Most of the current evidence of altered aggregation in presence of heterotypic interactions is in the form of morphology and kinetic data, but not structural insights. We unequivocally demonstrate that structural modulation of Aβ from its natural aggregation state does not necessarily have to arise from homotypic seeds: Aβ will undergo templated growth even when the seed fibrils are from a different isomer or mutant, and this is mediated through formation of heterotypic aggregates. Furthermore, while antiparallel intermediates for Iowa and Dutch mutants of Aβ have been identified^25,45^, equivalent structures for WT Aβ have only been speculated to exist but never experimentally identified. Our results essentially provide evidence that antiparallel fibrils of Aβ exist and can be formed through seeding from existing aggregates that share the same structural template. This also provides a rationalization of how such intermediates can be formed in general: not necessarily through homotypic seeds but through heterotypic interactions. This is particularly relevant when considering the structure of brain derived amyloid fibrils, which are typically generated through multiple generations of seeding from brain protein lysates^13,15,26,37^. However, it is challenging, if not impossible, to isolate only fibrillar seeds of specific Aβ isoforms from these lysates, which leads to the potential effect of multiple seeds from different isoforms of Aβ, of possibly different structures. Our results demonstrate that such seeds, that deviate from the commonly anticipated structures of Aβ, can lead to formation of daughter fibrils that also reflect these structural aberrations. As a result, when interpreting the structure of the daughter fibrils in such experiments, correlating them uniquely to specific polymorphs that prevail in amyloid plaques can be difficult. A key assumption underlying the characterization of brain derived fibrils is that they accurately represent the structural ensemble of aggregates that exist in the brain. Our results show that depending on the nature of the seeds, the daughter fibrils can have either morphology, structure or both that deviate from the seed. Recent NMR and cryo EM studies that have identified antiparallel structural motifs in brain derived fibrils^15,26^ underscore the significance of our findings and the importance of understanding structural propagation through homotypic and heterotypic seeding.

One important factor that must be considered in the context of the above results is that sequence homology of the seed and Aβ. While the seeds can act just as morphological templates of nucleation, structural propagation can be expected to be also facilitated through specific interactions between the sidechains of the seed fibrils and Aβ, which are maximized when their amino acid sequences overlap significantly, particularly of the amyloidogenic segments. In absence of either of these factors, i.e., when Aβ is aggregated in presence of an entirely different peptide in its monomeric/non-fibrillar form, it is reasonable to expect that no heterotypic or mixed aggregates will be formed. If this is true, this limits the biological significance of heterotypic aggregates, since they would require a very specific set of conditions to be fulfilled. To test this hypothesis, and whether the formation of heterotypic amyloid aggregate extends beyond the two interacting proteins with similar sequences, we chose to coaggregate two proteins with completely different sequences: ^13^C Aβ42 and α-synuclein. Both proteins were allowed to aggregate in a 1:1 mixture. AFM topographs, shown in Figure 5A, revealed the presence of fibrils after 24 h. The IR spectra obtained from different spatial locations on those fibrils can again be assigned to one of two subtypes (Figure 5B-D): one peak at ∼1590 cm^-1^ with a shoulder at ∼1660 cm^-1^ and the other peaks at ∼1660 cm^-1^ with a shoulder at ∼1590 cm^-1^. As per our observations on labeled Aβ42 detailed above, absorption at ∼1590 cm^-1^ confirms the presence of ^13^C Aβ42 in the aggregate whereas a peak at ∼1660 cm-1 denotes the presence of unlabeled peptide, i.e., α-synuclein, in addition to ^13^C Aβ42 in the fibrils. Furthermore, the ∼1590 cm^-1^ band is present in all the fibrillar spectra, indicating that α-synuclein does not aggregate independently and the resulting fibrils are all heterotypic in nature. This can be further verified by studying how pure α-synuclein aggregates in the same timeframe. The AFM topographs of α-synuclein aggregates after 24 h of incubation are shown in Figure S1, which clearly indicate a lack of fibrillar structure. The spectra acquired from these oligomeric and/or amorphous aggregates is also in agreement with the formation of heterotypic fibrils: pure α-synuclein spectra exhibit a peak at ∼1660 cm^-1^ and do not contain any significant intensity in the 1590 cm^-1^ spectral region (Figure S1E). To better understand the structural distributions in these heterotypic fibrils, their spectra, along with those of pure alpha synuclein, were deconvoluted by projecting onto the same MCR basis spectra as described above. The structural composition for both the heterotypic fibrils and α-synuclein, as obtained through the deconvolution, is shown in Figure 6. The α-synuclein aggregates are predominantly comprised of the non-β structural component, with a small contribution from β sheets. In contrast, the coaggregated fibrils have these components in addition to the labeled Aβ component, thus validating their heterotypic nature. These findings thus unequivocally demonstrate that even in the absence of a morphological template, Aβ can form mixed aggregates that incorporate other proteins with minimal amino acid sequence overlap. The two spectral subtypes mirror the relative variation of this component as seen in WT Aβ, suggesting that these fibrils do not particularly deviate structurally from WT. Interestingly, our findings somewhat deviate from expectations from previous coaggregation studies. The presence of other proteins during Aβ aggregation has been suggested to inhibit secondary nucleation, leading to prevention of fibril formation^33,50,51^. We observe that this does not apply to synuclein, and that Aβ can form heterotypic fibrils that incorporate the former. This in turn opens up further possibilities of how the Aβ aggregation pathway can be affected in presence of other brain proteins. To verify the generality of this finding and investigate whether Aβ42 can potentially form heterotypic fibrils if it is exposed to other brain proteins, we performed a similar co-aggregation experiment of ^13^C Aβ42 with protein lysate from a normal, non-AD brain. This represents a scenario where no pre-existing fibrillar seeds are present (Figure S1D). Furthermore, the amount of non-fibrillar Aβ is also expected to be minimal, as is expected for normal brain specimens and thus any interaction of Aβ42 with brain proteins is expected to be dominated by those which are non-Aβ in origin. We observe fibril formation after 24 h of incubation (Figure 5E), which again exhibit significant heterogeneity in spectral distribution and the amide I region of IR spectra obtained from different locations of these fibrils show underlying bands at 1600 cm^-1^, 1630 cm^-1^, 1640 cm^-1^ and 1680 cm^-1^ (Figure 5 F-H). This is markedly different from control ^13^C Aβ42 fibrils (Figure 1E) and the brain protein lysate spectra (Figure S1E) and thus clearly reveals the heterotypic nature of the fibrils. The weights obtained from MCR spectral deconvolution shows that the brain protein lysate in largely comprised of non-β sheet structures, with smaller populations of β sheets. In comparison, the fibrils have the additional β sheet band from ^13^C Aβ42, thus confirming their heterotypic nature. This further reinforces the possibility of Aβ interacting with non-homologous proteins when introduced to a heterogeneous mixture of multiple proteinaceous species. Of course, the relative abundances of these proteins and any Aβ fibrillar polymorphs that may additionally facilitate seeded growth will eventually dictate the complexity and heterogeneity of the overall aggregation process. Nonetheless, the results described here undoubtedly evidence the existence of alternative aggregation pathways in addition to usual homotypic seeding mechanism and emphasize the need to take into account the possibility of formation of fibrillar structures that are not necessarily from homotypic seeds when investigating brain derived aggregates.

**Figure 5.**
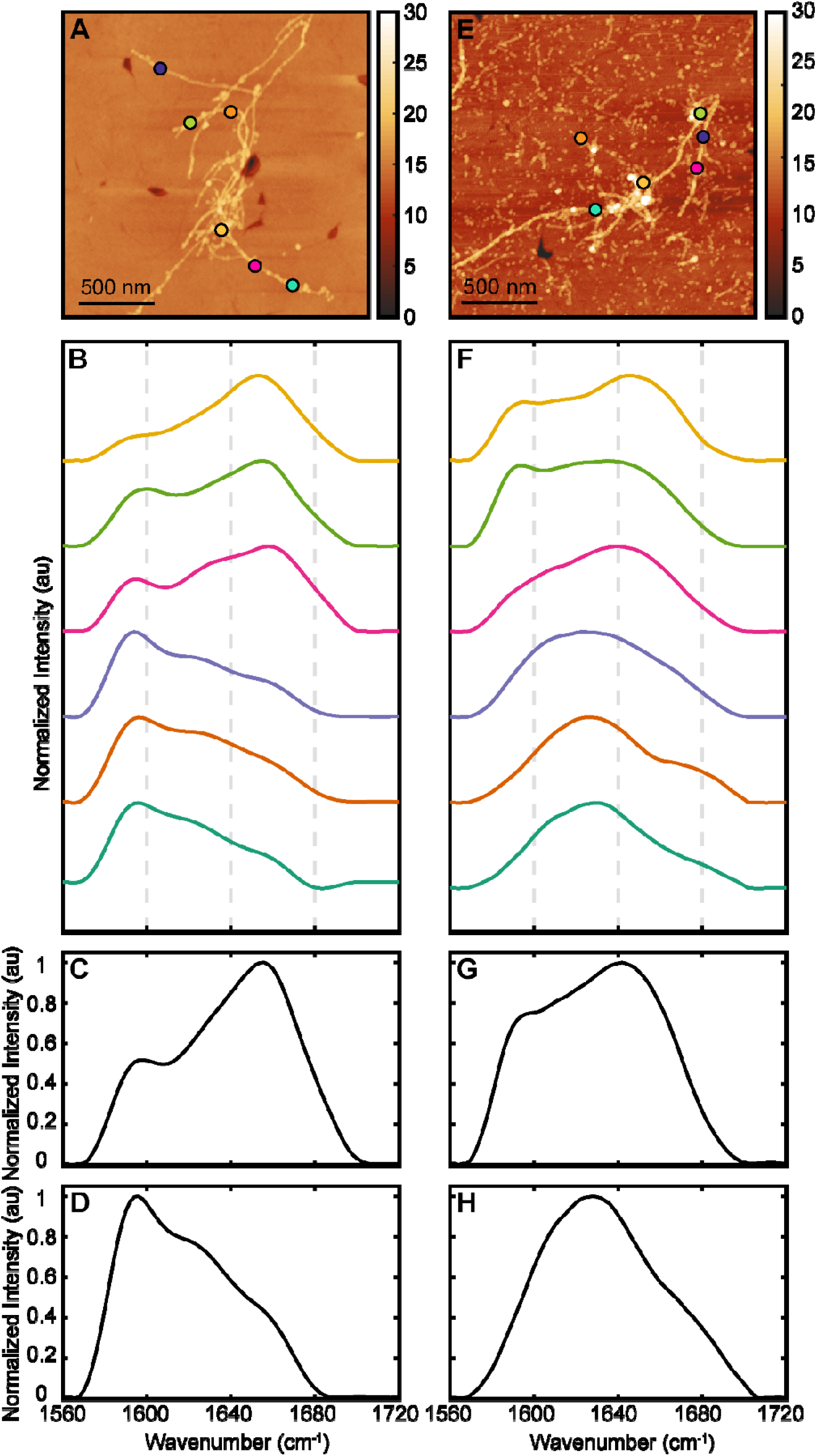
AFM-IR characterization of ^13^C-Aβ42, co-aggregated with alpha-synuclein (A-D) and with total brain protein lipid (E-H) in 10 mM phosphate buffer. (A,E) AFM topographic images of fibrils after 24 h of incubation. (B,F) Representative IR spectrum of co-aggregates recorded from the corresponding AFM images demonstrating the two spectral subtypes. (C,G) represents average spectra of subtype 1, while (G,H) denotes the average spectra of subtype 2.

**Figure 6.**
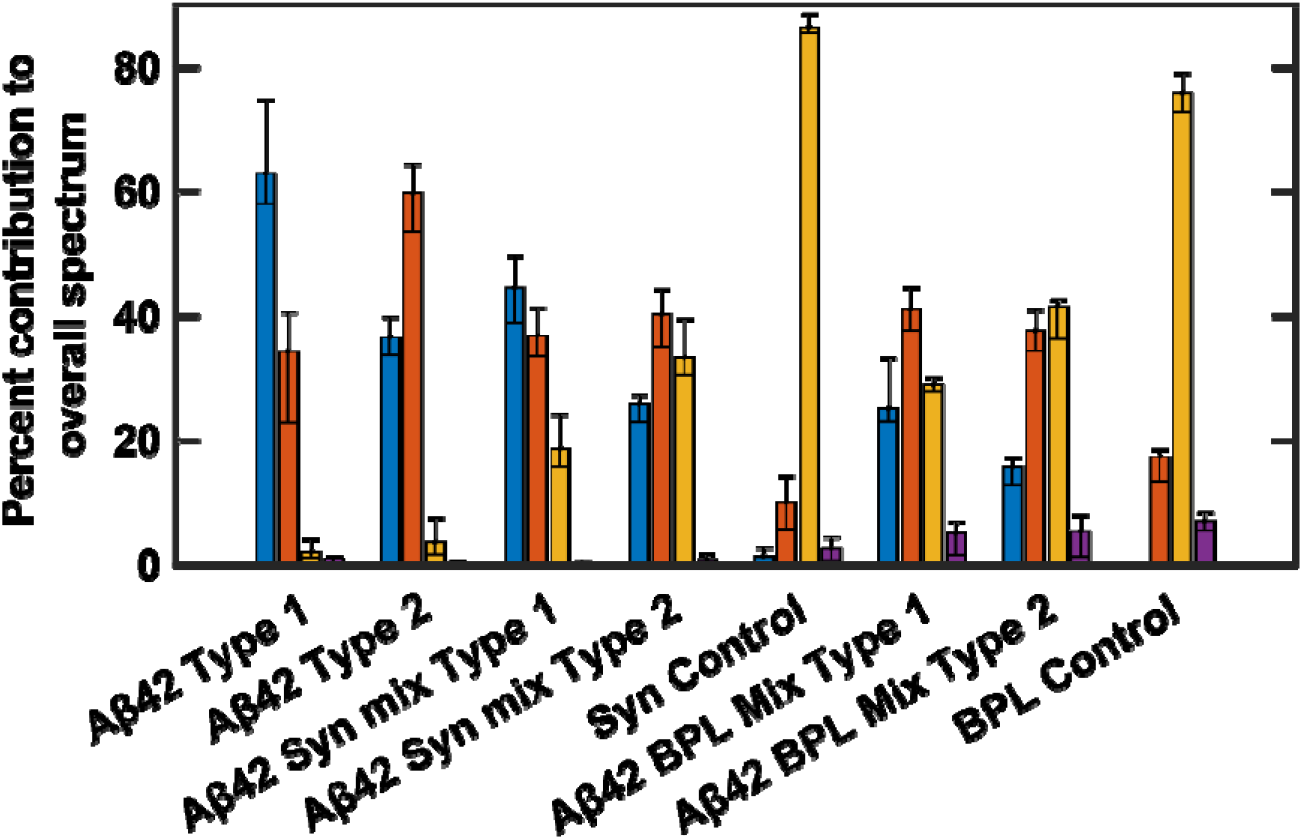
Secondary structure distribution of Aβ42 fibrils aggregated in presence of alpha-synuclein and brain protein lysate, as determined from MCR deconvolution. The composition of unseeded Aβ42 fibrils is also shown for comparison. The bars are color-matched to the spectral bands in Figure 4A. The error bars represent the interquartile range of the component weights.

## Conclusions

In summary, we demonstrate using spatially resolved nanoscale IR spectroscopy that heterotypic amyloid fibrils are formed when Aβ42 is allowed to aggregate in presence of either structurally different, preformed amyloid seeds or sequentially different monomeric protein. Furthermore, such heterotypic aggregates are also spontaneously generated when Aβ42 is exposed to total brain protein. The key insight that emerges from our findings is that the structure of amyloid fibrils can be significantly altered by heterotypic seeding, even if the structure of the seeds is different from native aggregates. One of the key challenges in NMR and cryo-EM is determining if seeded growth from brain protein extracts faithfully and accurately reproduces fibril structures found in the brain. In such contexts, heterotypic seeding is difficult to account for and often not considered. It is implicitly assumed that heterotypic seeding is an unlikely event and hence not a mechanism that can affect the fibril structure. We unequivocally demonstrate that a. heterotypic seeding of amyloid aggregates is possible, and b. can lead to formation of aggregates that are structurally distinct from the native Aβ fibrils. We further show that formation of heterotypic aggregates is not limited to different isoforms of Aβ only but can also spontaneously occur between Aβ and a non-homologous protein like alpha synuclein and in presence of brain protein extracts devoid of fibrillar seeds. Amyloid aggregates in the brain have a multiple number of proteins in addition to Aβ, and also various isomers of Aβ with different degrees of C and N terminal truncation. Smaller Aβ isoforms have been shown to form antiparallel aggregates in vitro, but it has been previously not understood how such aggregates would affect full-length Aβ. We show that seeding with antiparallel seeds leads to antiparallel fibrils of full length Aβ. Recent identification of antiparallel structures from the AD brain are consistent with our findings. This work shines light on the possibility of alternate heterotypic aggregation pathways of Aβ that are often not considered viable and thus promise to have far reaching impacts when assessing structure of brain derived amyloid aggregates.

## Data availability

All data required to evaluate the conclusions presented in the work are included in the manuscript and Supplementary Information. The datasets and any additional data are available from the corresponding author upon reasonable request.

## Author contributions

S.B. and D.B. conducted the experiments; S.B., D.B. and H.E. analyzed data; A.G. conceptualized and supervised this study; S. B., D. B., H.E. and A.G. wrote the manuscript.

## Acknowledgement

This work was supported by the National Institutes of Health (Award R35 GM138162 to A. G.). The content is solely the responsibility of the authors and does not necessarily represent the official views of the National Institutes of Health.

