## Supplementary Information for "Heterotypic Seeding Generates Mixed Amyloid Polymorphs"

**Materials and methods**

**Aggregation of ^13^C⁃labeled Aβ42**

^13^C-Aβ42 (rPeptide, USA) was first treated with 1,1,1,3,3,3-hexafluoroisopropanol (HFIP) to destroy any preformed aggregates. HFIP was completely evaporated at room temperature (24 °C), under vacuum. Concentration of ^13^C-Aβ42 was measured by UV spectrophotometer (Nanodrop, Thermo Scientific, USA), using an extinction coefficient of 1490 M^−1^ cm^−1^ at 280 nm, since Aβ42 has a single tyrosine residue. The aggregation of 100 μM ^13^C-Aβ42 was carried out in 10 mM phosphate buffer, pH 7.4 , at 37°C without any agitation.

**Aggregation of the seed:** **Aβ(16⁃22) and Dutch Aβ40**

Aβ(16⁃22) (rPeptide, USA) was first treated with 1,1,1,3,3,3-hexafluoroisopropanol (HFIP) to destroy any preformed aggregates. HFIP was completely evaporated at room temperature (24 °C), under vacuum. The aggregation of 500 μM Aβ(16⁃22) was carried out in 10 mM phosphate buffer, pH 7.4 , at 37°C without any agitation. The Aβ(16⁃22) seed was taken out after 6 hours of incubation as it already showed the presence of fibrils under AFM-IR, in the control experiment.

Similarly, the aggregation of 300 μM Dutch Aβ40 was carried out in 10 mM phosphate buffer, pH 7.4 , at 37°C without any agitation. Concentration of Dutch Aβ40 was measured by UV spectrophotometer (Nanodrop, Thermo Scientific, USA), using an extinction coefficient of 1490 M^−1^ cm^−1^ at 280 nm. The Dutch Aβ40 seed was taken out after 5 days of incubation.

**Aggregation of the α⁃synuclein**

α⁃synuclein (rPeptide, USA) was first treated with 1,1,1,3,3,3-hexafluoroisopropanol (HFIP) to destroy any preformed aggregates. HFIP was completely evaporated at room temperature (24 °C), under vacuum. Concentration of α⁃synuclein was measured by UV spectrophotometer (Nanodrop, Thermo Scientific, USA), using an extinction coefficient of 1490 M^−1^ cm^−1^ at 280 nm. The aggregation of 60 μM α⁃synuclein was carried out in 10 mM phosphate buffer, pH 7.4 , at 37°C without any agitation.

**Aggregation of ^13^C⁃labeled Aβ42 with Aβ(16⁃22)**

20 μM Aβ(16⁃22)seed was cross-seeded with 80 μM ^13^C-Aβ42 in 10 mM phosphate buffer, pH 7.4, for making the stock mixture. The aggregation was carried out at 37°C, without any agitation.

**Aggregation of ^13^C⁃labeled Aβ42 with Dutch Aβ40**

30 μM Dutch Aβ40 seed was cross-seeded with 60 μM ^13^C-Aβ42 in 10 mM phosphate buffer, pH 7.4, for making the stock mixture. The aggregation was carried out at 37°C, without any agitation.

**Coaggregation of ^13^C⁃labeled Aβ42 with α⁃synuclein**

60 μM α⁃synuclein was co-aggregated with 60 μM ^13^C-Aβ42 in 10 mM phosphate buffer, pH 7.4 for 24 h at 37°C without any agitation.

**Coaggregation of ^13^C⁃labeled Aβ42 with Total brain protein lysate (TBPL)**

500 µg/ml TBPL (BioChain Institute, CA) from normal human adult was co-aggregated with 100 µM of ^13^C- Aβ42 in 10 mM sodium phosphate buffer, pH 7.4 for 24 h at 37°C without any agitation. For control, 500 µg/ml TBPL was used.

**Sample preparation for AFM⁃IR experiment**

Samples were prepared by taking out aliquots from the co-aggregation mixture at 6 h and 24 h of incubation and depositing half-diluted solutions onto ultra flat gold substrates (Platypus Technologies, USA). 5 μl aliquot of the reaction mixture was incubated on the gold substrate for 5 min and then rinsed with 100 μl of Milli-Q water. Sample was dried with gentle stream of air and kept inside vacuum desiccator until imaging.

**AFM⁃IR experiment**

AFM-IR experiments were carried out by Bruker NanoIR3 instrument equipped with mid-IR quantum cascade laser (MIRcat, Daylight solutions). Experiments were performed at room temperature and relative humidity inside the instrument was kept low by continuous purging with dry air. Both AFM imaging and IR data collection was done in tapping mode with cantilevers having resonance frequency of 75±15 kHz and spring constant of 1-7 N/m. AFM scan rate were varied from 0.5 Hz to 1.0 Hz. First a high-resolution AFM image of the sample was recorded having multiple oligomers/fibrils in the scan area. Then AFM tip was placed on individual oligomer/fibril at random to obtain the IR spectra. Three different areas of sample were scanned where multiple oligomer/fibrils were probed for each time points to avoid any biasness or under sampling. IR spectral resolution was 2 cm^-1^. 128 coadditions at each point and 16 co-averages for each spectrum were applied.

**Data Analysis**

AFM images were processed by Gwyddion software. IR data were analyzed by using MATLAB software by applying (3, 7) Savitzky-Golay filter and a 5-point moving average filter. A baseline correction was applied for each spectrum.

**MCR-ALS**

MCR-ALS algorithm, implemented in MATLAB by Jaumot et al.,^1,2^ was used for spectral deconvolution. A total of 414 spectra (approximately 15-30 spectra per sample) was used for the deconvolution following the same exact spectral preprocessing highlighted previously in the manuscript. MCR-ALS essentially is a matrix factorization approach that determines the pure spectral responses (S) and their corresponding weights/concentrations (C) from a spectral dataset D as: D = C*T. The number of spectral components was chosen to be 4, which is consistent with the number of prominent peaks observed in the AFM-IR derivative spectra. Four Gaussian bands, centered at 1590 cm^–1^, 1630 cm^–1^, 1660 cm^–1^, and 1690 cm^–1^ were utilized as initial spectral estimates for the MCR-ALS algorithm. The corresponding weights/concentrations of each spectra were then divided by the sum of the concentrations to identify the percent contributions of each component.

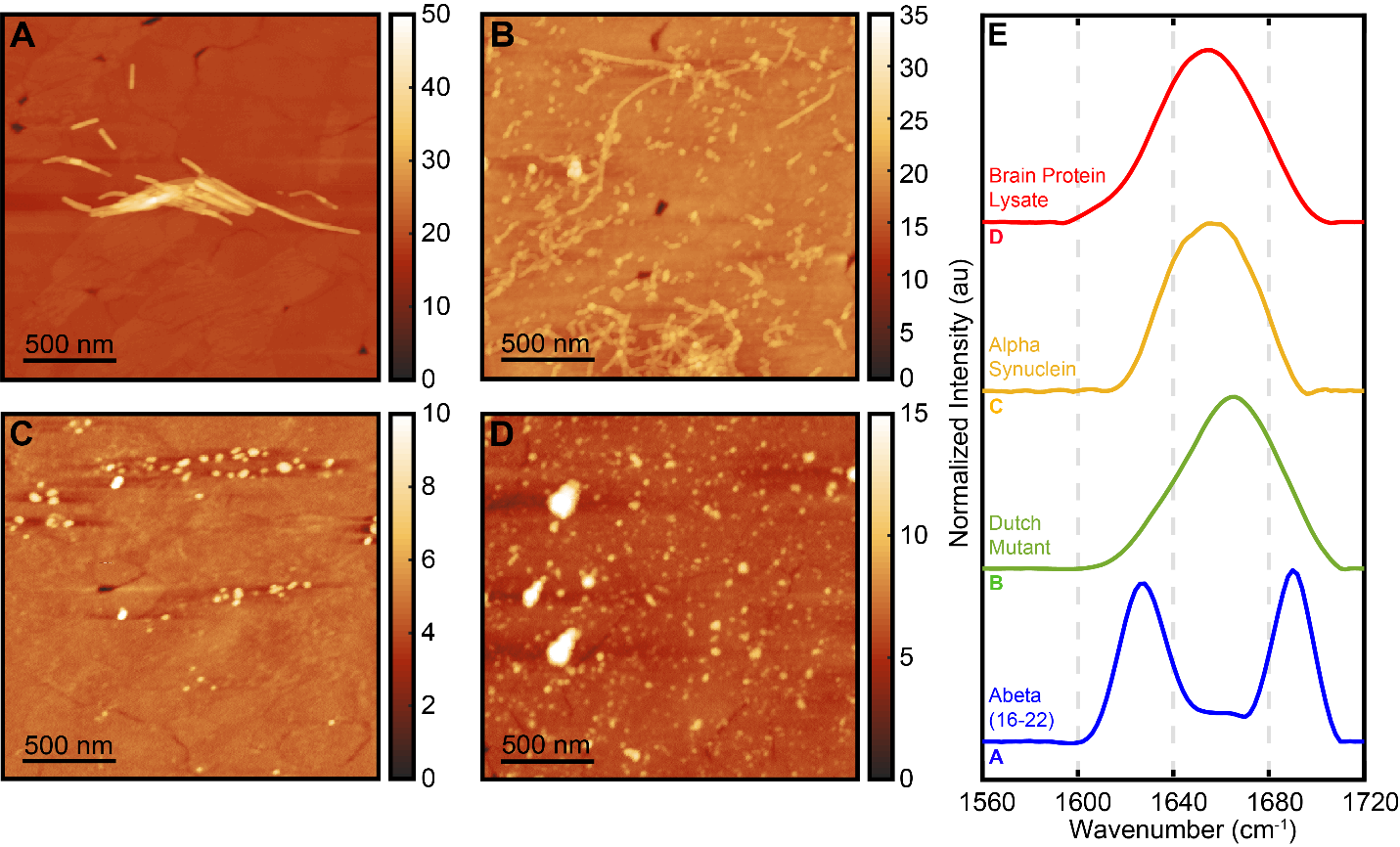

**Figure S1.** AFM topographic image of (A) Aβ(16-22) seed, (B) Dutch Aβ-40 seed, (C) α-synuclein control, and (D) TBPL control produced after 6 h, 5 days, 24 h and 0 h of incubation, respectively, at 37 ⁰C, without agitation. (E) Average IR spectrum from the corresponding AFM image.

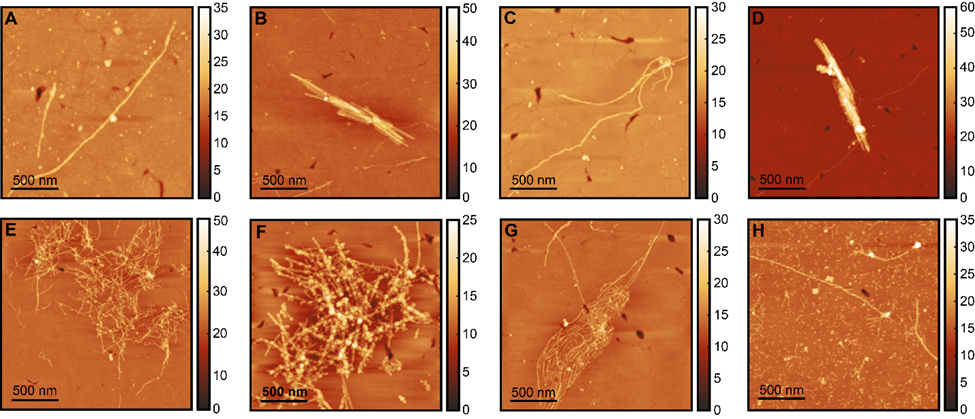

**Figure S2.** AFM topographs of fibrils generated from the cross-seeding mixture of ^13^C-Aβ42 with Aβ(16-22) seed after (A,B) 6 h and (C,D) 24 h of incubation, where (A,C) shows round fibrillar morphology and (B,D) represents flat fibrils. Bottom row shows AFM topographs of fibrils generated from the cross-seeding mixture of ^13^C-Aβ42 with Dutch Aβ-40 after (E) 6 h and (F) 24 h of incubation. (G,H) represents AFM topographs of fibrils produced from the co-aggregation of ^13^C-Aβ42 with α-synuclein and total brain protein lysate, respectively after 24 h of incubation.

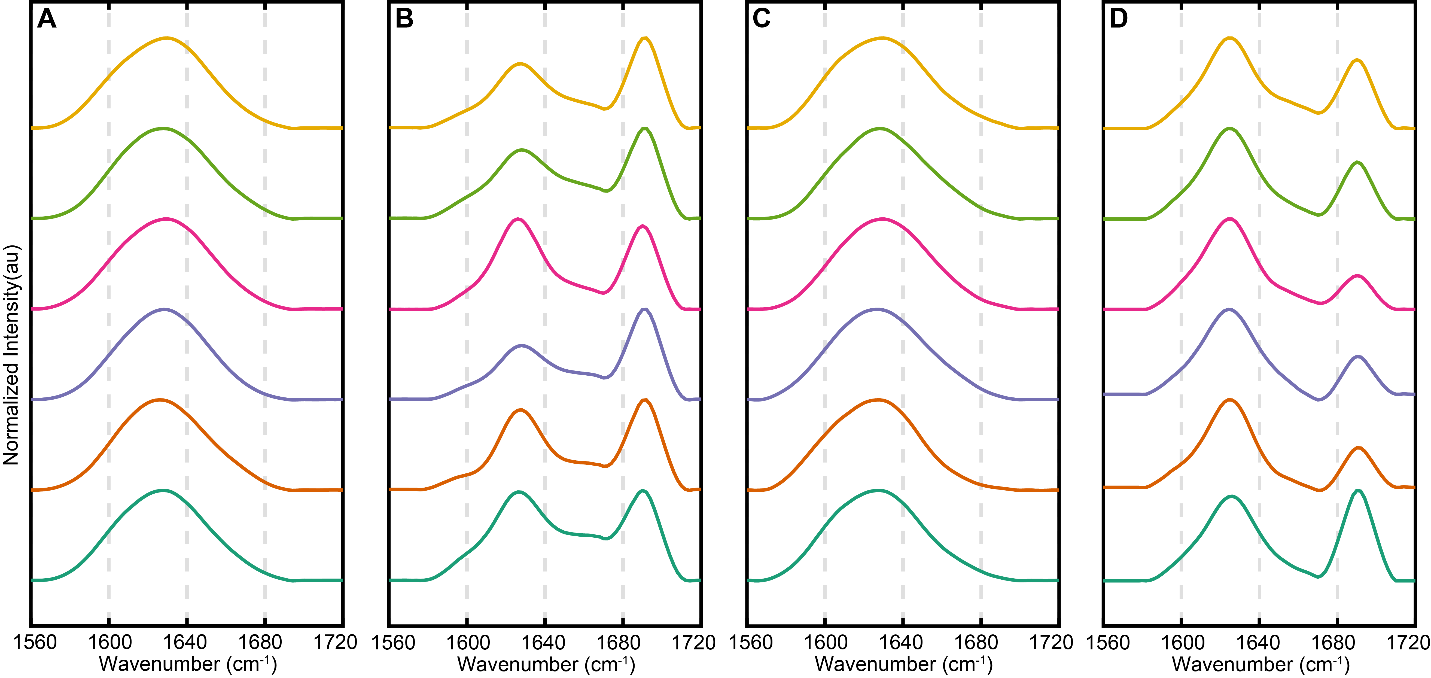

**Figure S3**. Representative IR spectra recorded from the fibrils, generated from the cross-seeding mixture of ^13^C-Aβ42 with Aβ(16-22) in 10 mM phosphate buffer, pH 7.4 after (A,B) 6 h and (C,D) 24 h of incubation. (A,C) represents first spectral subtype, arising from round fibrils, while (B,D) denotes the second spectral subtype, flat fibrils.

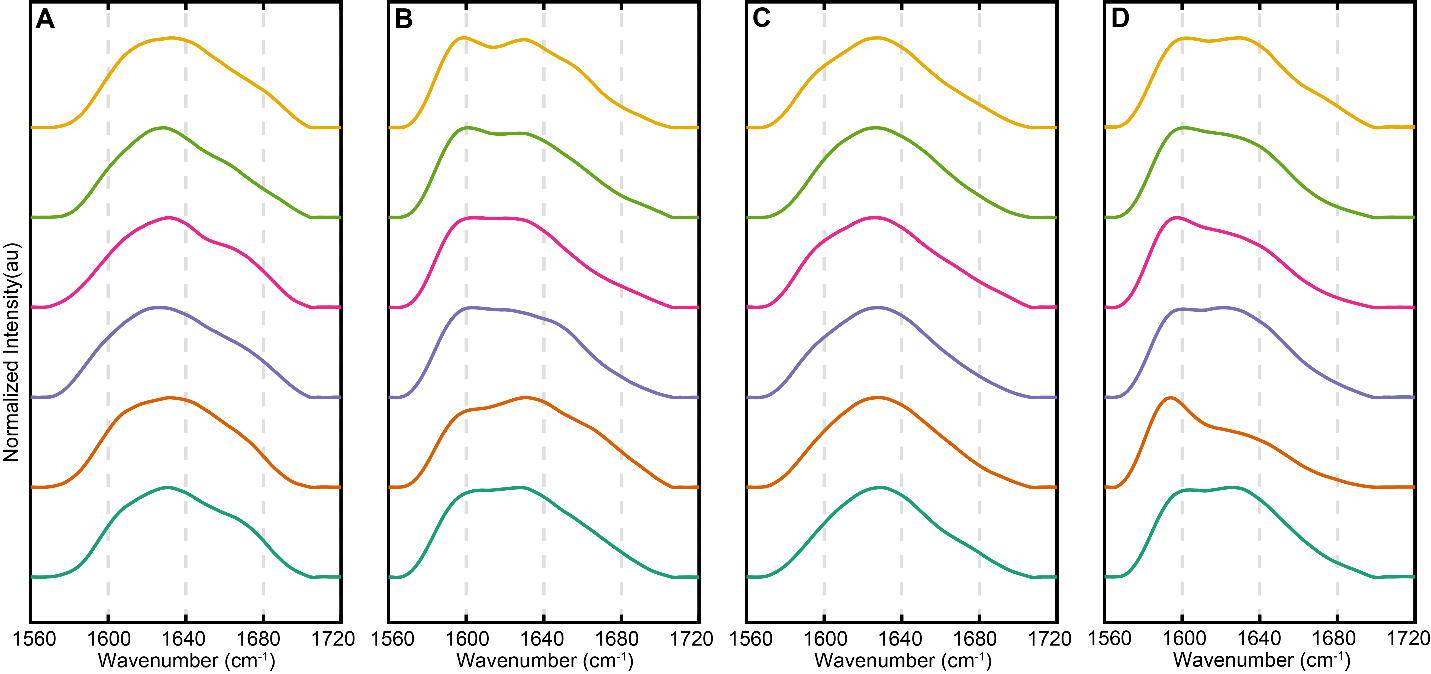

**Figure S4.** Representative IR spectra recorded from the fibrils generated from the mixture of ^13^C-Aβ42 cross-seeded with Dutch Aβ-40 in 10 mM phosphate buffer, pH 7.4 after (A,B) 6 h and (C,D) 24 h of incubation. (A,C) represents first spectral subtype, while (B,D) denotes the second spectral subtype.

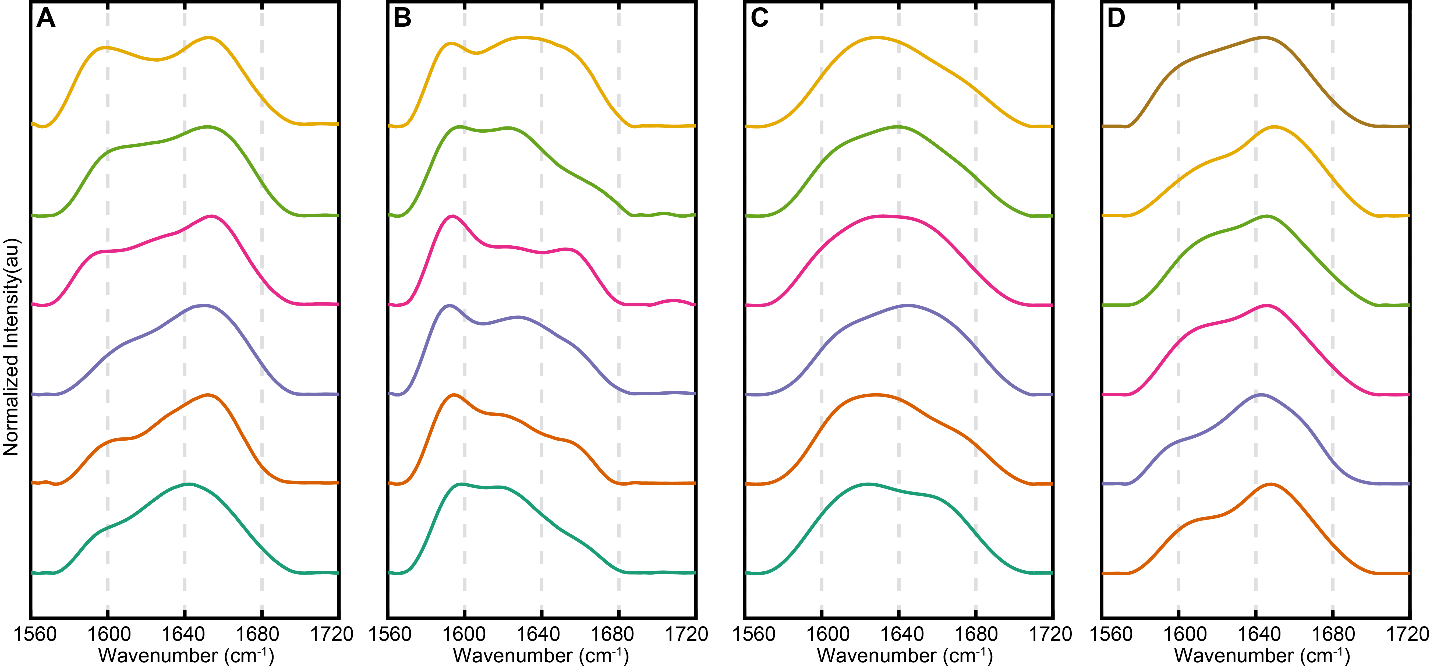

**Figure S5.** Representative IR spectra recorded from the fibrils generated from the co-aggregated mixture of ^13^C-Aβ42 with α-synuclein (A,B) and total brain protein lysate (C,D) in 10 mM phosphate buffer, pH 7.4, after 24 h of incubation. (A,C) represents first spectral subtype, while (B,D) denotes the second spectral subtype.
